# Genome-wide chromatin accessibility and transcriptome profiling show minimal epigenome changes and coordinated transcriptional dysregulation of hedgehog signaling in Danforth’s short tail mice

**DOI:** 10.1101/387977

**Authors:** Peter Orchard, James S. White, Peedikayil E. Thomas, Anna Mychalowych, Anya Kiseleva, John Hensley, Benjamin Allen, Stephen C. J. Parker, Catherine E. Keegan

## Abstract

Danforth’s short tail *(Sd)* mice provide an excellent model for investigating the underlying etiology of human caudal birth defects, which affect 1 in 10,000 live births. *Sd* animals exhibit aberrant axial skeleton, urogenital, and gastrointestinal development similar to human caudal malformation syndromes including urorectal septum malformation, caudal regression, VACTERL association, and persistent cloaca. Previous studies have shown that the *Sd* mutation results from an endogenous retroviral (ERV) insertion upstream of the *Ptf1a* gene resulting in its ectopic expression at E9.5. Though the genetic lesion has been determined, the resulting epigenomic and transcriptomic changes driving the phenotype have not been investigated. Here, we performed ATAC-seq experiments on isolated E9.5 tailbud tissue, which revealed minimal changes in chromatin accessibility in *Sd/Sd* mutant embryos. Interestingly, chromatin changes were localized to a small interval adjacent to the *Sd* ERV insertion overlapping a known *Ptf1a* enhancer region, which is conserved in mice and humans. Furthermore, mRNA-seq experiments revealed increased transcription of PTF1A target genes and, importantly, downregulation of hedgehog pathway genes. Reduced sonic hedgehog (SHH) signaling was confirmed by in situ hybridization and immunofluorescence suggesting that the *Sd* phenotype results, in part, from downregulated SHH signaling. Taken together, these data demonstrate substantial transcriptome changes in the *Sd* mouse, and indicate that the effect of the ERV insertion on *Ptf1a* expression may be mediated by increased chromatin accessibility at a conserved *Ptf1a* enhancer. We propose that human caudal dysgenesis disorders may result from dysregulation of hedgehog signaling pathways.

## Introduction

The semidominant Danforth’s short tail (Sd) mutation results in severe developmental abnormalities of the urogenital and gastrointestinal systems and the lower spine. These phenotypes overlap with caudal malformation syndromes in humans, including Currarino syndrome (OMIM #176450), urorectal septum malformation sequence (URSMS), VACTERL (Vertebral-Anal-Cardiac-Tracheo-Esophageal fistula-Renal-Limb anomalies) association (OMIM #192350), and caudal regression syndrome (CRS, OMIM #600145). Caudal birth defects affect 1:10,000 live births but their genetic etiology, intermediate molecular consequences to the epigenome and transcriptome, and resulting pathophysiology are largely unknown (1). The phenotypic overlap of the *Sd* mouse with multiple distinct human caudal dysgenesis disorders makes it an outstanding model to investigate the underlying etiology of human caudal dysgenesis.

Heterozygous (Sd/+) mice have a shortened tail, sacral vertebral anomalies, and frequent kidney anomalies but are viable and survive into adulthood (2). Homozygous *(Sd/Sd)* mice exhibit cessation of the vertebral column at the lumbar level and complete absence of the tail (2). Urogenital defects include agenesis or hypoplasia of the kidneys with occasional formation of the bladder and urethra and persistence of the cloaca. Gastrointestinal anomalies include imperforate anus. Homozygous animals are born in Mendelian ratios but are not viable beyond 24 hours (2). *Sd/Sd* and *Sd/+* embryos are visually distinguishable from wildtype (WT) embryos at E11 by caudal truncation and hemorrhagic lesions in the tailbud (3). Defects are apparent slightly earlier in *Sd/Sd* embryos. Prior to identification of the genetic lesion, differentiation of *Sd/Sd* and *Sd/+* embryos could only be made by the increased phenotypic severity in *Sd/Sd* mutants during gestation (4).

In 2013 our group and others identified the *Sd* mutation as an 8,528 bp insertion of an endogenous retroviral element (ERV) at a point 12,463 bp upstream of the *Pancreas specific transcription factor 1a (Ptf1a)* gene (5–7). The ERV insertion results in ectopic expression of *Ptf1a* at E8.5 in the notochord, lateral plate mesoderm and tail mesenchyme that persists into the caudal notochord, tailbud mesenchyme, mesonephros, and hindgut of E9.5 embryos (6). The critical role of PTF1A in causing the *Sd* phenotype was demonstrated by knockout of *Ptf1a* genomic sequences followed by replacement of *Ptf1a* coding sequences, which resulted in attenuation and recapitulation of the *Sd* phenotype, respectively (7).

Ptf1a is a basic helix-loop-helix (bHLH) transcription factor that is expressed in the pancreas and in neuronal progenitors in the cerebellum, hindbrain, neural tube and retina (8–11). Null mutations in *Ptf1a* in humans and mouse result in pancreatic and cerebellar agenesis (12, 13). PTF1A interacts with an E-box protein and RBPJ to form the trimeric PTF1 complex. PTF1 binds a bipartite cognate site including a canonical E-box motif (CANNTG) and TC-box (TGGGAAA) (14). PTF1 drives initial pancreatic organogenesis as well as the specification of GABAergic neurons in the neural tube (13, 15, 16). In the pancreas, PTF1 subsequently drives differentiation of mature acinar cells (9). Masui et al. characterized a *Ptf1a* autoregulatory enhancer region, which contains two PTF1 motifs located 14.8 and 13.5kb upstream of *Ptf1a* (17). Both enhancers drive *LacZ* expression in the dorsal and ventral pancreas and the neural tube (NT) (17). Neural tube-specific enhancers have been identified 12.4kb downstream of *Ptf1a* coding sequences (18, 19). Two long non-coding RNAs (lncRNA’s), *Gm13344* and *Gm13336,* have been annotated in the genomic region surrounding *Ptf1a* in the mouse. Interestingly, the transcription start site (TSS) for *Gm13344* overlaps the region corresponding to the *Ptf1a* 13.5kb upstream autoregulatory enhancer, while *Gm13336* overlaps *Ptf1a* coding sequences (17).

Our previous studies of the *Sd* mutation implicated ectopic *Ptf1a* expression in the *Sd* phenotype. The genome-wide epigenome and transcriptome alterations induced by the *Sd* mutation are largely uncharacterized. To address this issue, we measured chromatin accessibility using ATAC-seq and transcriptional profiles using mRNA-seq on E9.5 WT and *Sd/Sd* tailbuds (20–22). We found minimal changes to the genome-wide landscape of chromatin accessibility - only one peak met our genome-wide 5% False Discovery Rate (FDR) threshold. Strikingly, the peak is adjacent to the *Sd* insertion site, which is located at the *Ptf1a* 13.5kb upstream autoregulatory enhancer that overlaps with the *Gm13344* TSS. This peak is more open in *Sd* mice compared to WT. The mRNA-seq results confirmed upregulation of *Ptf1a* and *Gm13344* in *Sd* mutants and identified significantly (<5% FDR) reduced expression of genes within the Hedgehog signaling pathway. In-situ hybridization confirmed downregulation of *Shh* as well as other notochordal markers. Further, dysregulation of caudal neural tube patterning was observed in *Sd/Sd* embryos, consistent with *Shh* downregulation. Finally, we observed increased apoptosis in the tailbud and caudal somites. Our findings suggest that ectopic *Ptf1a* expression, potentially driven by increased chromatin accessibility at an upstream enhancer, induces a cascade of transcriptional dysregulation of multiple genes in the Hedgehog signaling pathway. We propose that ectopic *Ptf1a* ultimately leads to downregulation of Shh signaling, degeneration of the notochord, and increased apoptosis in the developing caudal region.

## Results

### ATAC-seq reveals localized changes in chromatin accessibility near the Sd insertion

Transcription factor (TF) binding and chromatin accessibility are strongly coupled. Closed chromatin and inaccessible DNA may prevent a TF from binding a target site; on the other hand, cooperative binding of TFs, or binding of certain ‘pioneer factors’, may increase or maintain chromatin accessibility. Therefore, the over- or under-expression of TFs may alter chromatin accessibility (23–26). *Ptf1a* is strongly overexpressed in *Sd* mice and is a direct regulator of other TFs. The *Sd* insertion is located adjacent to a *Ptf1a* autoregulatory enhancer region. In order to quantify changes in chromatin accessibility in *Sd* mutant embryos, we performed ATAC-seq on E9.5 WT and *Sd/Sd* tailbuds. We analyzed four WT and four *Sd/Sd* ATAC-seq libraries, pooling tailbuds from two embryos for each library to ensure sufficient input material. ATAC-seq peak calling identified 26,620 reproducible peaks (see Methods). A differential peak analysis uncovered little change in ATAC-seq signal genome wide, revealing only one significantly differential peak (unadjusted p-value 2.2×10^−7^; adjusted p-value 5.8×10^−3^; 5% FDR threshold for the analysis) (27). This peak corresponds to the promoter of *Gm13344,* a lncRNA located 13.7kb upstream of *Ptf1a* (Fig. 1A, B). The peak called at the *Ptf1a* promoter region was not significantly different between WT and *Sd/Sd* E9.5 embryos at FDR 5%; however, the peak is skewed towards being more open in the WT (unadjusted p-value 4.5×10^−4^; adjusted p-value 0.61; Fig. 1B). This suggests that while there are few genome-wide changes in chromatin accessibility in the Danforth mouse, local chromatin accessibility near *Ptf1a* and the *Sd* insertion site is altered.

**Fig. 1:**
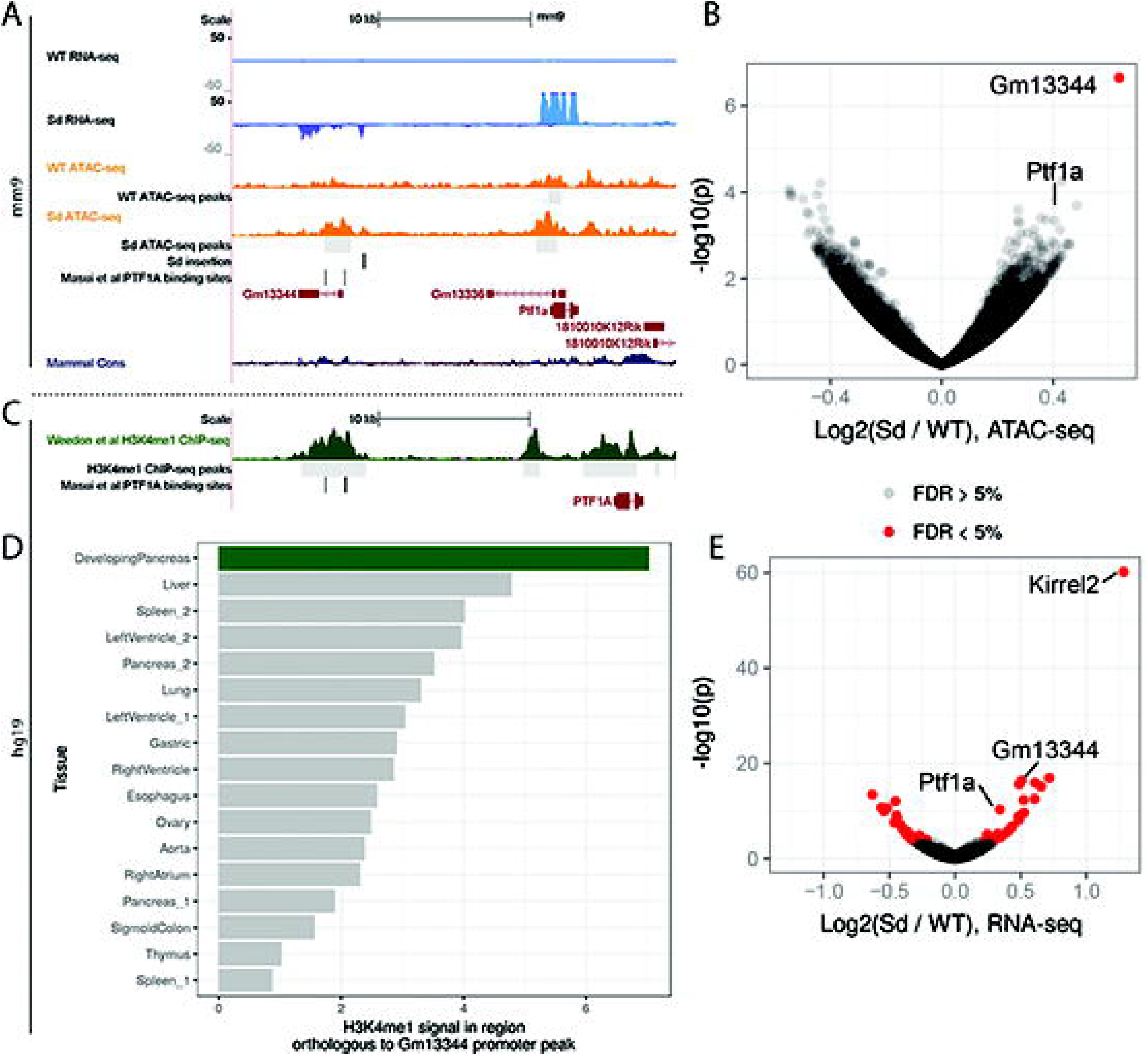
Chromatin accessibility and transcriptome changes occur at the *Ptf1a* locus of *Sd/Sd* mutant embryos. (A) (A) UCSC Genome Browser (Kent et al 2002) displaying ATAC-seq (orange) and RNA-seq signal (blue) near the *Sd* insertion, *Ptf1a,* and *Gm13344.* (B) Volcano plot for ATAC-seq data. The gene represented by the nearest TSS for the significantly differential peak is indicated, as is the peak representing the *Ptf1a* promoter. Red dots denote significance at FDR <5%. (C) Weedon et al. ChIP-seq data (30), indicating a peak above the human region orthologous to the *Gm13344* promoter. (D) The ChIP-seq signal for H3K4me1 in developing pancreas (from Weedon et al) is greater than the signal in other tissues/at other time points. (E) Volcano plot for RNA-seq data.

### Gm13344 promoter is orthologous to a human pancreatic developmental enhancer

The mouse lncRNA *Gm13344* is not present in standard human genome annotations such as Gencode and RefSeq. In a previous study, expression of a transgene containing the *Sd* ERV and *Gm13344* in mice did not lead to the development of a short tail, indicating that *Gm13344* overexpression alone is unlikely to cause the Danforth phenotype (7). LncRNAs are generally not conserved across species and may evolve from unstable transcription of regulatory elements (28, 29). Masui et al previously found that in the human genome, PTF1A binds an enhancer element ~15kb upstream of the *PTF1A* gene containing two PTF1A motif-matches that are conserved in mouse (Fig. 1A) (17). Interestingly, the orthologous mouse Ptf1a binding sites occur near the *Gm13344* promoter, suggesting that the mouse *Gm13344* lncRNA promoter may be functionally comparable to the human enhancer. In order to determine whether this human enhancer is active during pancreatic development, we uniformly processed and analyzed publicly available ChIP-seq data for H3K4me1, a histone mark that colocalizes with enhancers, from developing human pancreatic cells and other control tissues (control tissues downloaded from Roadmap Epigenomics) (30–32). Developmental pancreas H3K4me1 data show a clear H3K4me1 peak at the enhancer, consistent with the enhancer being active during human pancreatic development. (Fig. 1C). Notably, H3K4me1 ChIP-seq data from other tissues and time points display lower signal in the region orthologous to the mouse *Gm13344* promoter (Fig. 1D). Collectively, these results suggest that the Sd-insertion-mediated increase in chromatin accessibility at the *Gm13344* promoter acts as an enhancer for *Ptf1a* in E9.5 mice, and transcription of the lncRNA is at least partially a byproduct of this regulatory activity.

### mRNA-seq reveals misexpression of Ptf1a target genes

To determine which genes show evidence of misregulation in *Sd* mutant embryos at E9.5, we performed mRNA-seq on three WT and three *Sd/Sd* E9.5 tailbud libraries (samples were pooled as necessary for sufficient input material to each library; see Methods and S3 Table). The WT tailbud transcriptome can be found in the supplemental data (S1 Table). We performed differential gene expression analysis and found 49 genes to be significantly differentially expressed (5% FDR; Fig. 1E, Fig. 2). Several of the differential genes discovered by mRNA-seq recapitulate previous findings. As expected, *Ptf1a* showed almost no expression in WT tailbuds but strong expression in *Sd/Sd* tailbuds (Fig.1A, Fig. 2). Similarly, *Gm13344,* the lncRNA upstream of *Ptf1a,* showed almost no expression in WT tailbuds but strong expression in the *Sd* mutant samples, consistent with previous reports (Fig. 1A, Fig. 2) (7). Carboxypeptidase A1 *(Cpa1),* previously found to be underexpressed in E10.5 *Ptf1a* knockout (KO) mice, is overexpressed in *Sd* mutant tailbuds (Fig. 2) (33). We found *Foxa1* to be downregulated in *Sd* mutant samples (Fig. 2), consistent with the upregulation of *Foxa1* in *Ptf1a* knockout mice (33). *Foxa2* also trended towards downregulation in *Sd* mice, although the adjusted p-value did not reach significance (unadjusted p-value 2.0 x 10^-4^, adjusted p-value 0.066) (Fig. 2). Two of the most strongly upregulated genes in *Sd/Sd* tailbuds were *Kirrel2* and *Nphs1* (Fig. 1E, Fig. 2), which share a bidirectional promoter to which *Ptf1a* is known to bind at later time points in neural tube and pancreas (34). *Aplp1,* positioned directly downstream of *Kirrel2,* is similarly upregulated in *Sd* mutant samples (Fig. 2). mRNA-seq also revealed upregulation of *Sox8* in *Sd* mutant tailbuds (Fig. 2). SOX8 is a High Mobility Group (HMG) transcription factor that functions redundantly with SOX9 and SOX10 in neural crest development and its expression precedes that of SOX9 and SOX10 (35). No significant differences in the caudally expressed genes *Wnt3a, Cyp26a1,* and *Cdx2* were identified in *Sd* mutant samples in our mRNA-seq data (Fig. 2).

**Fig. 2:**
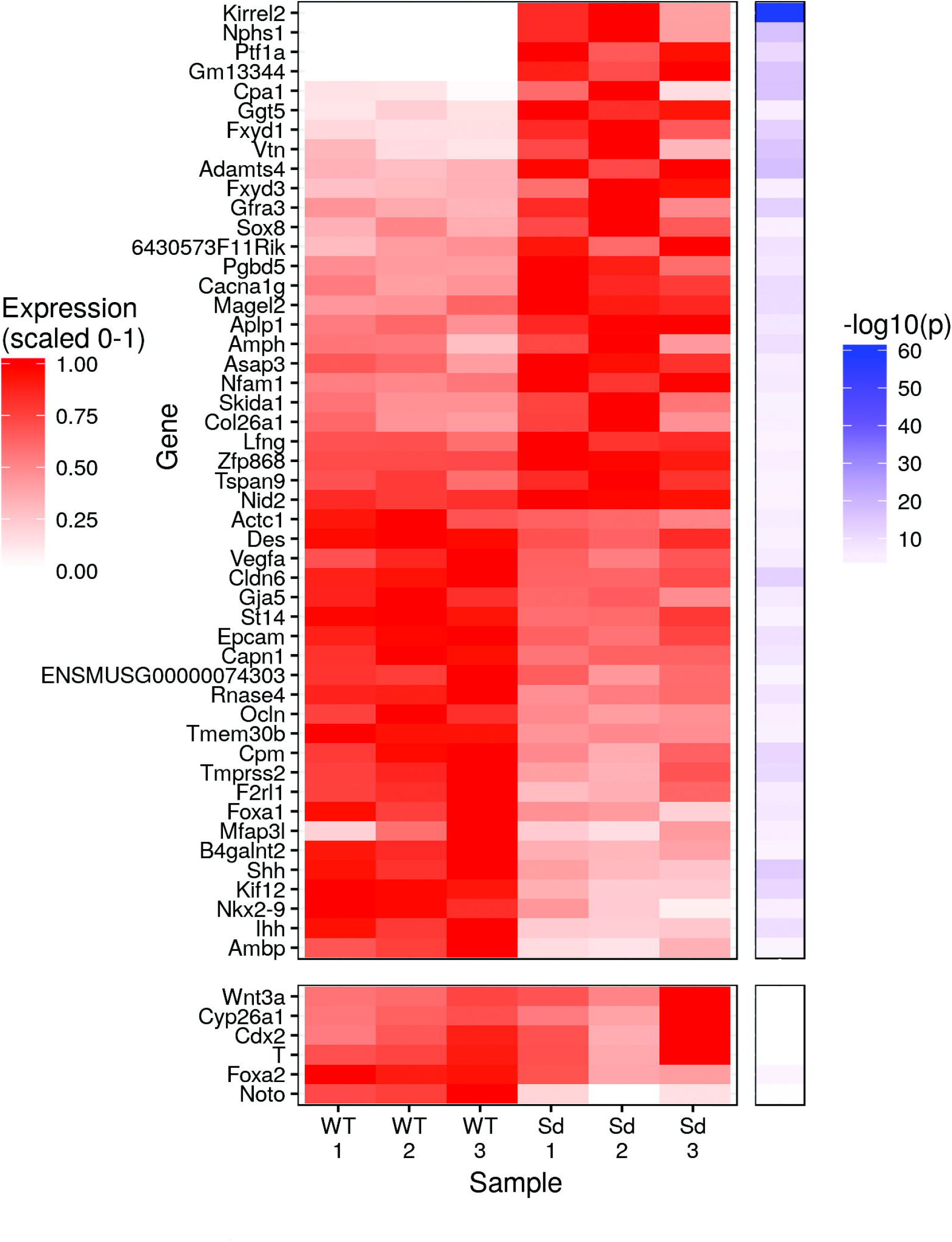
Ectopic *Ptf1a* expression drives global transcriptomic changes within *Sd/Sd* embryo tailbud. Heatmap displays normalized expression and -log10(p) for differentially expressed genes and other genes of interest.

### Whole mount in situ hybridization analysis validates mRNA-seq expression changes

To confirm expression changes in *Sd* mutant tailbuds identified by mRNA-seq analysis, we used whole mount in situ hybridization in E9.5 embryos. These studies confirmed ectopic expression of *Ptf1a* in the tailbud of E9.5 *Sd/+* and *Sd/Sd* embryos (S1 Fig). The ectopic *Ptf1a* expression was stronger in homozygous mutants compared to *Sd/+* embryos, and appeared highest in the presomitic mesoderm with lower levels in the notochord in the anterior thoracic region. In addition, we found increased expression of *Gm13344* in the tailbud of *Sd/+* and *Sd/Sd* embryos (S1 Fig). We also found upregulated expression of *Kirrel2* (Fig. 3A-C) in *Sd/+* and *Sd/Sd* mutants, as identified by mRNA-seq. *Foxa1* expression was clearly reduced in *Sd/Sd* embryos (Fig. 3D-F), in accord with our mRNA-seq data. *Foxa2* expression was also reduced in the tailbud by whole mount in situ hybridization (Fig. 3G-I). Although the unadjusted p-value for *Foxa2* was significant in the mRNA-seq analysis, it did not survive genome-wide multiple testing correction using 5% FDR. This could be due to the relatively small proportion of *Foxa2*-expressing cells in the tailbud (36). We also examined expression of the T-box transcription factor brachyury (T) in the tailbud of *Sd/Sd* embryos. *T* is expressed in notochord-derived cells and is required for notochord development, mesoderm differentiation and development of structures posterior to the forelimb level in mouse embryos (37–39). We found that *T* expression was unaffected in the tailbud of *Sd* mutant embryos, although reduced expression was observed in the more anterior part of the notochord of *Sd/Sd* embryos (Fig. 3J-L), which correlates with our mRNA-seq data (Fig. 2). *Noto,* a Not family homeobox gene, is expressed in the wild type mouse tailbud and is essential for proper caudal notochord function (36). We did not detect *Noto* expression in the tailbud of *Sd/Sd* embryos, and expression was decreased in the *Sd/+* embryos (Fig. 3M-O), which could indicate suppression of its expression, since *T* expression indicates the presence of the caudal notochord in *Sd/Sd* embryos at this timepoint.

**Fig. 3:**
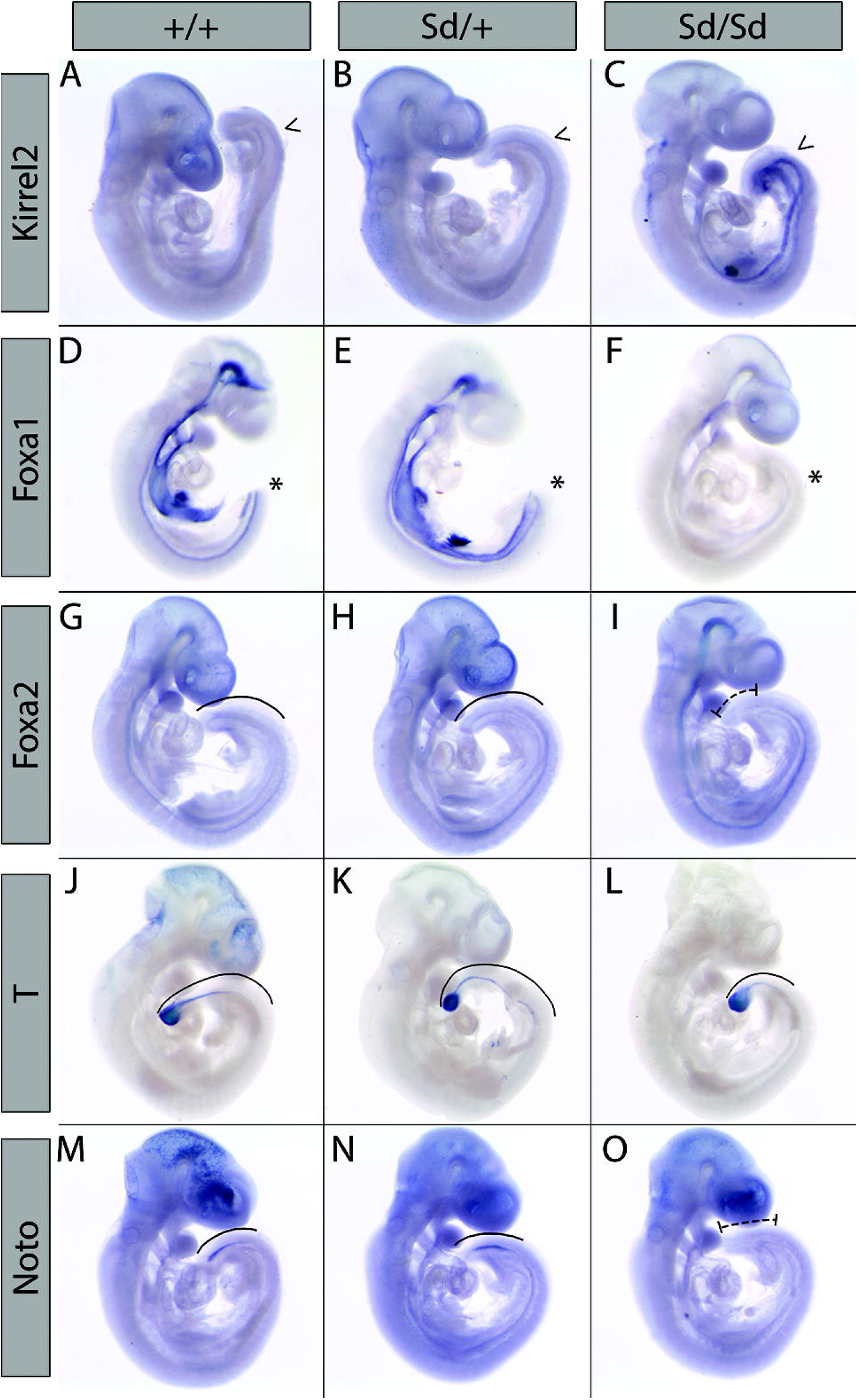
Whole mount in situ hybridization confirms transcriptomic changes in *Sd/Sd* embryos. Whole mount in situ hybridization with antisense probes for *Kirrel2* (A-C), *Foxa1* (D-F), *Foxa2* (G-I), *T* (J-L), and *Noto* (M-O) in E9.5 WT, *Sd/+,* and *Sd/Sd* embryos. Arrow head and asterisk indicate increased and decreased, respectively, expression patterns in the tailbud of *Sd/Sd* embryos. Solid lines indicate staining within the tailbud (G,H,J,K,M,N), dashed lines (I,O) indicate extent of reduced staining in Sd/Sd tailbuds. N > 3 for each genotype per probe.

### Pathway analysis shows dysregulated hedgehog signaling

To determine which developmental signaling pathways are disrupted in *Sd* mutant tailbuds, we performed a KEGG pathway enrichment analysis using the p-values of all genes included in the differential gene expression analysis. Several pathways related to cellular adhesion and migration show significant enrichment for disrupted gene expression (Fig. 4A). Interestingly, the only developmental pathway to show significant enrichment for altered gene expression was the Hedgehog signaling pathway (Fig. 4A). Sonic Hedgehog (Shh) and Indian Hedgehog *(Ihh)* are two of the most strongly downregulated genes in *Sd* mutant samples (Fig. 2A), and in total seven genes annotated to hedgehog signaling show an unadjusted p-value for differential expression of less than 0.05 *(Shh* (p=4.43 × 10^−14^), *Ihh* (p=6.62 × 10^−10^), *Gli1* (p=0.018), *Gli3* (p=0.019), *Ptch1* (p=0.021), *Ptch2* (p=0.002), *Bmp4* (p=0.006)) with changes in the direction expected for downregulation of this pathway (S2 Table). Whole mount in situ hybridization confirmed that *Shh* expression was absent in the tailbud of *Sd/Sd* embryos at E9.5, and its expression decreased in a reciprocal manner to the ectopic expression of *Ptf1a* in the tailbud (Fig. 4B-D). NKX2.9 is a homeobox transcription factor expressed in close proximity to SHH in neural tube structures along the entire neuraxis and is regulated by SHH (40, 41). *Nkx2.9* expression was significantly reduced in *Sd/Sd* embryos by mRNA-seq and confirmed by whole mount in-situ hybridization (Fig. 2A and Fig. 4E-G). *Gli1,* a major downstream effector of SHH signaling, was reduced by LacZ staining in mice from a cross between *Sd/+* and *Gli1-LacZ* reporter mice (Fig. 4H, I) in accordance with SHH downregulation.

**Fig. 4:**
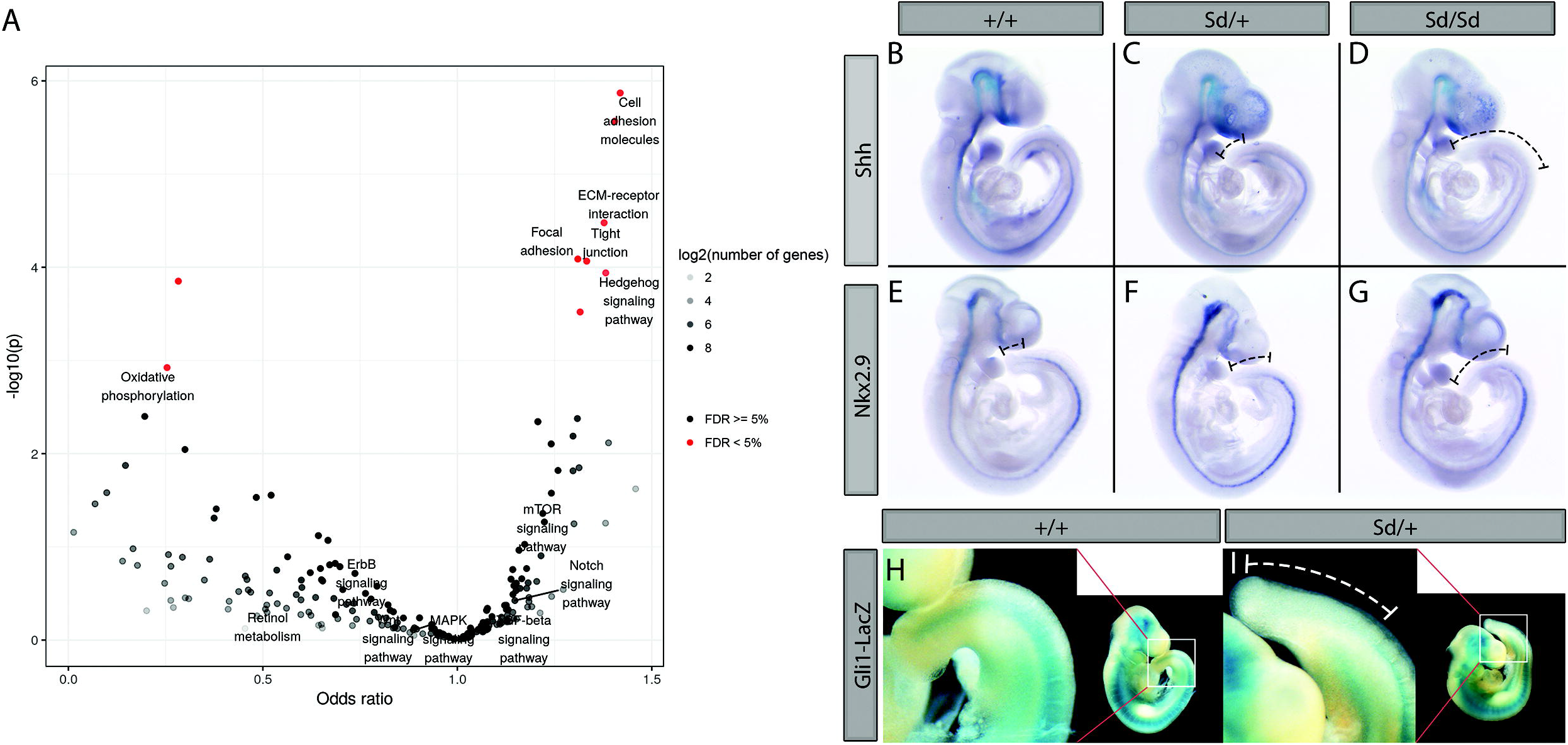
KEGG pathway analysis of *Sd/Sd* embryos identifies dysregulated SHH signaling. (A) Volcano plot for enrichment of disrupted gene expression in KEGG pathways. Notable developmental signaling pathways are labeled. Hedgehog signaling is the only developmental signaling pathway showing an enrichment for differential gene expression. (B-D) Whole mount in situ hybridization with *Shh* and *Nkx2.9* (E-G) antisense probes in E9.5 WT, *Sd/+,* and *Sd/Sd* embryos. (H,I) X-gal staining of E10.5 WT and *Sd/+* embryos from a cross between *Sd/+* mice and *Gli1-lacZ* reporter mice. N > 3 for each genotype of stained embryos.

### The absence of Shh results in aberrant caudal neural tube patterning in Sd/Sd embryos

We hypothesized that dysregulation of SHH signaling in developing *Sd* mutant embryos would result in aberrant neural tube patterning. To test this hypothesis, we conducted immunofluorescence experiments on E10.5 WT, *Sd/+* and *Sd/Sd* mice. Embryos were sectioned in the transverse plane through the neural tube and notochord at the level of both the forelimb and hindlimb bud. Staining with an antibody against SHH revealed aberrant notochordal morphology in *Sd/+* embryos and absence of the notochord in *Sd/Sd* embryos (Fig. 5A-F) Floorplate expression of SHH was maintained throughout *Sd/+* embryos; however, it was lost at the hindlimb level of *Sd/Sd* embryos. Floorplate-derived SHH is induced following FOXA2 initiation in the floorplate by notochordal-derived SHH (42–46). Thus, SHH loss in the floorplate is likely due to the downregulation of FOXA2 in the floorplate (Fig. 5G-L, E’, F’). Ventralization of the neural tube relies on SHH signals; thus, loss of the V3 domain marker NKX2.2 in the *Sd/Sd* hindlimb neural tube is consistent with SHH loss (Fig. 5M-R, G’, H’) (47). Interestingly, the more widely expressed marker NKX6.1 is expressed but ventrally constricted in *Sd/Sd* mutant embryos (Fig. 5S-X). Consistent with the loss of SHH signaling from the floorplate and notochord, the dorsally restricted transcription factor PAX6 exhibits ventral expansion at the hindlimb level of *Sd/Sd* embryos (Fig. 5Y-D’) Reduced expression of ventral markers as well as expansion of dorsal neural tube markers at the hindlimb level in the neural tube of *Sd/Sd* embryos indicates the notochord of *Sd/Sd* mutant embryos degenerates before correct patterning of the presumptive floorplate can be established.

**Fig. 5:**
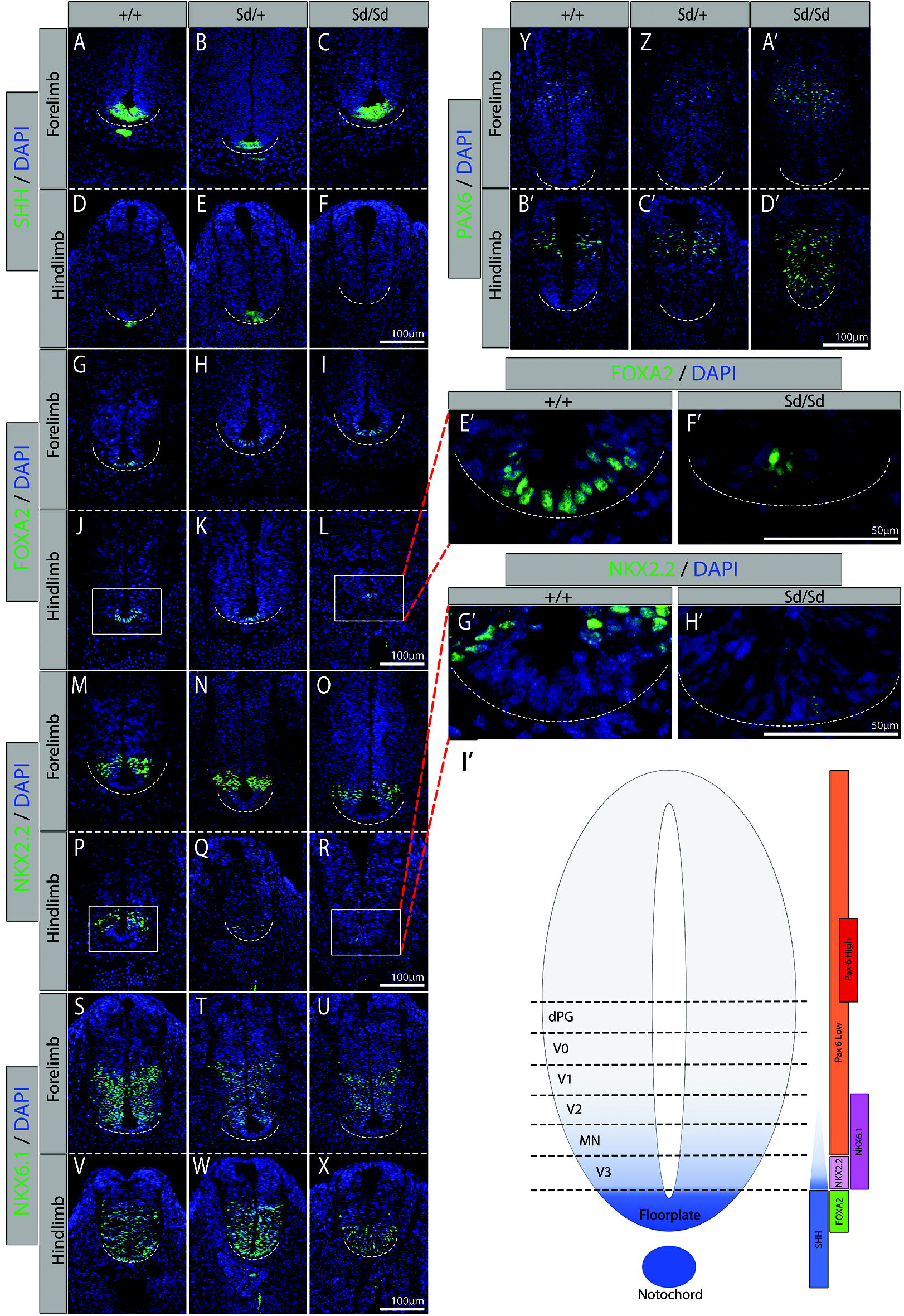
Immunofluorescence staining confirms dysregulated SHH expression in *Sd/Sd* embryos and identifies aberrant neural tube patterning. Immunofluorescence studies of SHH expression (green) in WT, (A,D) *Sd/+,* (B,E) and *Sd/Sd* (C,F) embryos as well as downstream targets FOXA2, (G-L) NKX2.2, (M-R) NKX6.1 (S-X) in transverse forelimb (top pane) and hindlimb (bottom pane) sections. Dorsal marker PAX6 (Y-D’) indicates the competing WNT/ BMP gradient. Enlarged FOXA2 (E’,F’) and NKX2.2(G’,H’) floorplate images of WT(J,P) and *Sd/Sd* (L,R) images respectively. WT expression domains of these transcription factors (I’). All sections co-stained with DAPI for histological reference (blue). Dashed line indicates ventral border of neural tube. N > 3 for each genotype per antibody.

### Sd mutant embryos exhibit increased apoptosis in the tailbud

Previous studies have revealed hemorrhaging within the regressing tailbud of E12.5 *Sd* mutant embryos; however, degeneration of the tailbud in *Sd* mutant embryos has not been closely investigated (4). No visible phenotypic differences were apparent at E9.5. However, we found that hematomas are clearly evident in *Sd/Sd* embryos at E10.5, which likely give way to severe hemorrhaging as development progresses (Fig. 6A-C). Further investigation of the tailbud degeneration using immunofluorescence with an antibody to cleaved caspase-3 revealed increased apoptosis in *Sd/Sd* tailbud tissues relative to WT and *Sd/+* littermates (Fig. 6D-F). Increased apoptosis could be a result of loss of hedgehog patterning from the notochord in *Sd/Sd* embryos (48–52).

**Fig. 6:**
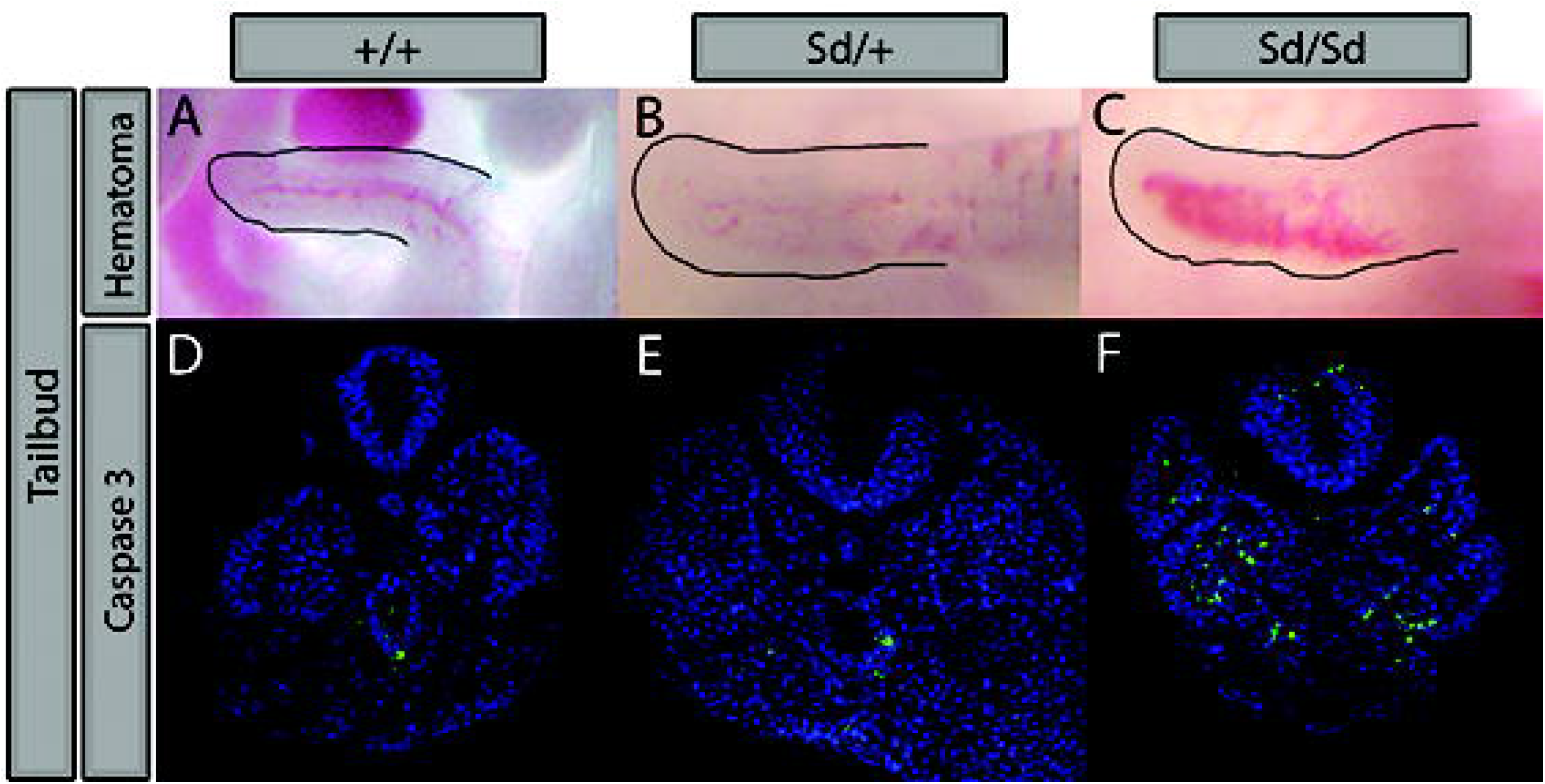
*Sd/Sd* tailbuds exhibit gross hematomas and increased apoptosis. WT(A), *Sd/+(B)* and Sd/Sd(C) tailbuds imaged immediately following dissection in PBS, hematoma is evident in *Sd/Sd* embryo tailbud. Increased apoptosis was observed by immunofluorescence staining for Caspase 3 in *Sd/Sd* tailbud (F) compared to WT(D) and *Sd/+(E)* tissues. N > 3 for each genotype.

## Discussion

Our group and others previously identified the *Sd* mutation as an ERV insertion upstream of *Ptf1a* that leads to its ectopic expression during caudal development, resulting in the *Sd* mutant phenotype (5–7). The underlying mechanisms that lead to upregulation of *Ptf1a* and the downstream developmental pathways dysregulated by ectopic *Ptf1a* expression were not previously determined. In the current study, we used ATAC-seq to analyze genome-wide chromatin accessibility and mRNA-seq to gain additional insight into these underlying mechanisms.

Our ATAC-seq data showed a significant change in chromatin accessibility near the *Ptf1a* genomic locus, mapping to the promoter region of *Gm13344,* a lncRNA that has been annotated in the mouse genome. This lncRNA overlaps with a previously characterized autoregulatory enhancer of *Ptf1a* (17). According to our mRNA-seq data, *Gm13344* is transcribed and overexpressed in *Sd/Sd* tailbuds relative to WT tailbuds. Previous studies also detected overexpression of *Gm13344* in *Sd* mutant embryos, although a transgene containing the *Sd*

ERV and *Gm13344* without *Ptf1a* did not recapitulate the *Sd* phenotype, while a transgene containing the *Sd* ERV and *Ptf1a* did recapitulate some, but not all, of the *Sd* phenotype (7). We previously generated a transgene containing 31.9 kb of genomic sequence surrounding *Ptf1a* with and without the *Sd* ERV, which did not recapitulate the *Sd* phenotype (5). This transgene contained part of the *Gm13344* sequence, including the PTF1A binding sites. The discrepancy between these transgene experiments suggests that the genomic regulatory context is critical for appropriate localization of ectopic *Ptf1a* expression and may have been strongly influenced by positional integration effects of the transgenes. Thus, the exact effects of *Gm13344* on the *Sd* phenotype are not known.

We found no evidence of an annotated human lncRNA that is orthologous to the mouse *Gm13344* lncRNA. However, the promoter region is conserved in humans and maps to a region that contains high H3K4me1 signal in developing pancreas relative to other tissues, including adult pancreas (30). This finding suggests that the enhancer properties of the *Gm13344* orthologous promoter sequence in humans are conserved. Our results suggest that the genomic sequence corresponding to the *Gm13344* promoter in mouse functions as an enhancer, consistent with previous studies, and the resulting RNA produced from *Gm13344* is a byproduct of this enhancer activity, as has been shown for other developmentally regulated genes (17, 29, 53). In addition, we speculate that it is the combination of transcription factors binding to the *Sd* ERV and the *Gm13344* enhancer region that is required for the tissue and temporal specificity of ectopic *Ptf1a* expression in *Sd* mutant embryos. A corollary of this hypothesis is that ectopic *Ptf1a* feeds into this autoregulatory loop to further increase ectopic *Ptf1a* expression.

Our transcriptome analysis revealed 49 genes with significant differential expression at 5% FDR. As expected, *Ptf1a* and *Gm13344* were significantly upregulated, as were known PTF1A downstream targets, including *Kirrel2, Nphs1,* and *Cpa1. Kirrel2* and *Nphs1* share a bidirectional promoter, which contains known PTF1A binding sites. They were originally identified as components of the interdigitating podocyte foot processes of the glomerular filter but are also expressed in the developing nervous system (54). Expression of both *Kirrel2* and *Nphs1* are lost in *Ptf1a* null mice, while forced expression of *Ptf1a* in the cerebral cortex induced ectopic expression of both genes (55). Therefore, the transcriptomic changes we identified were widespread but characterized by expected changes due to upregulation of *Ptf1a* expression.

One of the most striking findings revealed by our mRNA-seq analysis was that the hedgehog signaling pathway is altered in *Sd* mutant embryos. Both *Shh* and *Ihh* were significantly downregulated, and KEGG pathway enrichment analysis revealed the Hedgehog pathway as significantly dysregulated compared to other known developmental signaling pathways. These findings were confirmed by in situ hybridization analysis with *Shh,* as well as a cross of *Sd/+* mice to the *Shh* reporter strain *Gli1-LacZ,* which revealed decreased β-galactosidase expression in *Sd/+* tailbuds. We also confirmed reduced expression of *Nkx2.9* and *Foxa2,* which are regulated by SHH along the neural axis (56).

Our mRNA-seq data did not demonstrate significant differences in expression of *Cdx2, T, Wnt3a,* and *Cyp26a1,* and the KEGG pathway enrichment analysis did not identify the *Wnt* or Retinoic acid signaling pathways as being significantly dysregulated. Because *T* is expressed in notochord-derived cells and is required for notochord development, we performed in situ hybridization analysis for *T* expression (39, 57, 58). While we did observe reduced *T* expression in the anterior notochord in *Sd/Sd* embryos, expression of *T* in the tailbud was unchanged. Although our data conflict with a previous study, our analysis of tailbud tissue and use of more specific methodology (RNA-seq and in situ hybridization compared to qRT-PCR in RNA from whole embryos) could explain this apparent discrepancy (7). We observed increased expression of *Lfng* in *Sd/Sd* tailbuds by mRNA-seq. *Lfng* is a downstream effector of *Notch1* signaling. Although RBPJ is an integral component of the PTF1 complex, its expression was unchanged in *Sd* mutant tailbuds (unadjusted p-value = 0.56), and the *Notch* signaling pathway was not found to be dysregulated by KEGG pathway analysis. Since different E-box proteins are used as the third component of PTF1 complex depending on the context, it is difficult to predict which E-box proteins might be dysregulated in Sd mutant embryos. In addition to T, we also examined expression of *Noto* as a marker of notochord expression (36, 59, 60). Not surprisingly, we found that *Noto* expression was absent in the caudal notochord of *Sd/Sd* embryos. It is possible that the absence of *Noto* expression is due to degeneration of notochord cells in this region. However, the persistence of *T* expression in the caudal notochord suggests that *Noto* expression is suppressed either directly or indirectly by ectopic *Ptf1a.*

FOXA2 induces *Shh* expression in both notochordal and floorplate tissues, and importantly, floorplate-derived SHH is sufficient to pattern the neural tube (42–46, 61). Immunofluorescence staining through the transverse plane of forelimb and hindlimb level neural tube revealed normal expression of SHH in the floorplate of *Sd/+* embryos, which is sufficient to correctly maintain the expression domains of FOXA2, NKX2.2, NKX6.1, and PAX6 in the E10.5 neural tube, despite aberrant notochord morphology and reduced SHH signaling from this organizing tissue.

Similarly, in *Sd/Sd* mutant embryos, normal expression of SHH from the floorplate is sufficient to pattern the neural tube; however, only at the forelimb level. In *Sd/Sd* embryos, SHH is completely absent from the presumptive floorplate at the hindlimb level and as a result, dorsal neural tube transcription factor domains of FOXA2, NKX2.2, and NKX6.1 are reduced, and expression of the ventrally constricted transcription factor PAX6 is expanded dorsally. Taken together, these experiments demonstrate that dysregulated SHH signaling adversely affects patterning of the neural tube in *Sd/Sd* embryos. If dysregulated SHH signaling drives the *Sd* phenotype, then the increased phenotypic severity of *Sd/Sd* embryos could result from the degeneration of the notochord before redundancy between the floorplate and notochord has been established in the caudal *Sd/Sd* embryo.

Numerous published studies in mouse models have demonstrated that caudal and urogenital malformations resembling the malformations observed in *Sd* mutants can be secondary to disrupted SHH signaling. The *Shh* knockout mouse (62) exhibits abnormal axial structures including the floorplate, ventral neural tube and sclerotome. Additional mouse models with disrupted SHH signaling have been demonstrated to have phenotypes of anorectal, renal, and vertebral malformations that are observed in *Sd* mutant mice, resembling VACTERL association (63–65). Runck et al (66) studied cloacal development in *Shh* knockout mice and in human patients with cloacal malformations and found striking similarities between the two. Knockout of *Shh* in the notochord or floorplate using either *ShhCreERT2* or *Foxa2CreERT2* mice exhibit a phenotype that is strikingly similar to *Sd* mutant mice only in the notochord knockout (67). Knockout of both *Foxa1* and *Foxa2* in the notochord results in severe defects in formation of the axial skeleton and dorsal-ventral patterning of the neural tube secondary to reduced *Shh* expression (68), also phenocopying *Sd* mutant mice. Finally, *in vivo* knockdown of *T* in the notochord results in reduced *Shh* expression in the notochord and also phenocopies the *Sd* mutant phenotype (58). Thus, the caudal phenotype observed in *Sd* mutant mice is consistent with previous reports of disrupted SHH signaling in the notochord.

Collectively, our results show that the *Sd* insertion leads to widespread transcriptional dysregulation of the hedgehog developmental signaling pathway. The effect of the *Sd* insertion may be mediated by a nearby conserved *Ptf1a* autoregulatory enhancer, which shows increased chromatin accessibility in *Sd* mutant mice. More generally, our results suggest a cascade of molecular events where a single non-coding regulatory mutation propagates to perturb a critical signaling pathway. Further, our findings suggest that regulatory mutations in hedgehog signaling pathway genes could explain some human caudal malformations without known genetic etiologies. Future studies of viable mouse models, such as the *Sd* mouse, with alterations in SHH signaling, will be critical to dissecting the timing and impact of key molecular events and will help to further our understanding of human caudal malformations.

## Materials and methods

### Animals

All animals were housed in environmentally controlled conditions with 14 hour light and 10 hour dark cycles with food and water provided *ad libitum.* All Protocols were approved by the Institutional Animal Care & Use Committee (IACUC) at the University of Michigan and comply with policies, standards, and guidelines set by the State of Michigan and the United States Government. *Sd/+* animals were maintained on an outbred *CD-1* background (Charles River Laboratories, Wilmington, MA).

### Timed pregnancies

Matings for timed embryo isolation were set up using standard animal husbandry techniques. Noon on the day of vaginal plug observation was considered E0.5. Embryos were isolated at E9.5-E10.5. Yolk sac DNA was isolated via the HotSHOT extraction method for genotyping. Genotyping was performed as previously described (5).

For RNA-seq and ATAC-seq, embryo tailbuds were dissected just prior to the terminal somite, snap frozen in liquid nitrogen and stored at −80C.

### mRNA-seq experiments

RNA was collected from the tailbuds using a dounce homogenizer and RNeasy mini extraction kit (Qiagen). First, samples were placed into a dounce homogenizer with 350 uL of RLT buffer and homogenizing with 5 strokes. The extraction procedure was followed as specified by the kit, including the optional DNAse treatment. Samples were eluted in 35 uL of RNAse free water. Libraries were prepared at the University of Michigan DNA Sequencing Core using the Illumina TruSeq stranded mRNA-seq kit.

### ATAC-seq experiments

All procedures were done at 4C to minimize degradation of nuclei, and following the previously published protocol except as noted here (20). Each tailbud was disrupted in 200 uL of NIB with 0.5% Triton-X buffer by pipetting up and down. Then the samples were spun, supernatant removed and was followed by a second wash with RSB buffer. Each pellet of nuclei was then transposed using a home-made Tn5 enzyme (21, 69). After 30 minutes of incubation at 37C, each sample was cleaned up with MinElute PCR purification kit (Qiagen).

### Sequencing data

Libraries were multiplexed and sequenced on an Illumina HiSeq 2500 instrument. All raw and processed data have been deposited to GEO under the accession: GSE108804.

### ATAC-seq data processing

Adapters were trimmed using cta (v. 0.1.2). Reads were aligned to mm9 using bwa mem (v. 0.7.15-r1140; flags: -M (70). Picard MarkDuplicates (v. 2.8.1) was used for duplicate removal (options: VALIDATION_STRINGENCY=LENIENT) and samtools (v. 1.3.1) was used to filter for autosomal, properly-paired and mapped read pairs with mapping quality >= 30 (samtools view – b -h -f 3 -F 4 -F 8 -F 256 -F 1024 -F 2048 -q 30) (71). Peak calling was performed using MAC2 callpeak (v. 2.1.1.20160309; options: --nomodel --broad --shift −100 --extsize 200 --keep-dup all) (72). Peaks were filtered against a blacklist (downloaded from http://mitra.stanford.edu/kundaje/akundaje/release/blacklists/mm9-mouse/mm9-blacklist.bed.gz). For figures displaying ATAC-seq coverage, we normalized the signal to account for differences in library size. This was done by dividing each line in the *treat_pileup* bedgraph files generated during peak calling with MACS2 by the number of tags in the treatment (‘total tags in treatment’ as output by MACS2) and multiplying by 10 million. The bedgraph files were then converted to bigwig format using bedGraphToBigWig (v. 4).

### Differential peak calling

The list of reproducible peaks for downstream analysis was generated by taking the union of all 5% FDR broad peaks across all eight samples, and keeping the union peaks that overlapped with 5% FDR peak calls from at least two of the eight ATAC-seq libraries. Differential peak calling was performed using DESeq2 (v. 1.14.1) (27). Read counts for each sample and each peak were collected using bedtools’ coverageBed (v. 2.26.0; coverageBed -counts) (73). The number of somites in each sample (S3 Table) was used as a covariate (in the case that the sample consisted of pooled embryos, the number of somites for the sample was set to the mean number of somites in the embryos). Differential peaks were called at 5% FDR.

### RNA-seq processing and differential gene expression analysis

Reads were aligned to mm9 (Gencode vM1 comprehensive gene annotation) using the STAR splice-aware aligner (v. 2.5.2b; --outSAMUnmapped Within KeepPairs) (74, 75). Aligned reads were filtered to autosomal reads with mapping quality 255 (samtools view -b -h -f 3 -F 4 -F 8 -F 256 -F 2048 -q 255). We used QoRTs (v. 1.0.7) to gather read counts per sample per gene for differential gene expression analysis (76). Differential gene expression analysis was performed using DESeq2 (5% FDR). The number of somites in each sample (S3 Table) was used as a covariate. For figures displaying RNA-seq coverage, we normalized the signal to account for differences in library size. To do this, the filtered RNA-seq bam files were converted to wiggle format using QoRTs’ bamToWiggle (with options --negativeReverseStrand --stranded -- sizefactor XXX), where XXX represents the corresponding sample’s size factor (from QoRTs function get.size.factors).

### GO enrichment

GO enrichment analysis using the differentially expressed genes was performed using RNA-Enrich (with the KEGG database; code downloaded from supplemental materials of (77)). Gene sets with less than 5 genes or more than 500 genes were excluded from the analysis; otherwise, default parameters were used.

### ChIP-seq processing

The accession numbers of the data we used from Weedon et al. and Roadmap Epigenomics are listed in S4 Table (30, 32). We aligned this data using bwa aln (default parameters). All reads were trimmed to 36 bps (using fastx_trimmer from FASTX toolkit v 0.0.14; http://hannonlab.cshl.edu/fastx_toolkit) to prevent read length from confounding comparisons. Picard MarkDuplicates was used for duplicate removal (options: VALIDATION_STRINGENCY=LENIENT) and samtools was used to filter to autosomal, properly-paired and mapped read pairs with mapping quality >= 30 (samtools view -b -h -F 4 -F 256 -F 1024 -F 2048 -q 30). Peak calling was performed using MAC2 callpeak (options: --broad --keep-dup all). Peaks were filtered against hg19 blacklists (downloaded from http://hgdownload.cse.ucsc.edu/goldenPath/hg19/encodeDCC/wgEncodeMapability/wgEncodeDacMapabilityConsensusExcludable.bed.gz and http://hgdownload.cse.ucsc.edu/goldenPath/hg19/encodeDCC/wgEncodeMapability/wgEncode_DukeMapabilityRegionsExcludable.bed.gz). We normalized the signal for each experiment to 10M reads to prevent sequencing depth from confounding comparisons. Quality control was performed using phantompeakqualtools (v.2.0); all samples included in the analysis had RSC > 0.8 and NSC > 1.05. (78, 79). One of the two Weedon et al. H3K4me1 samples (ERR361008) appeared as a QC outlier (NSC = 1.31, RSC = 2.67) relative to other samples (mean NSC = 1.10, mean RSC = 1.47) and was therefore excluded (30).

To determine the human sequence orthologous to the *Gm13344* promoter peak, we used bnMapper with an mm9 to hg19 chain file from UCSC (downloaded from http://hgdownload-test.cse.ucsc.edu/goldenPath/mm9/liftOver/mm9ToHg19.over.chain.gz) to map the *Gm13344* promoter peak onto the human genome (80). The resulting closely-spaced stretches of human orthologous sequence were then merged together (bedtools merge –d 30) in order to eliminate small gaps between them, and the signal for this region in each tissue’s H3K4me1 ChIP-seq experiment was used for the comparison in (Fig. 1C).

### Reproducibility of computational analyses

We have created a GitHub repo that has all the code and steps necessary to reproduce the analyses in this work. The repo is located at: https://github.com/ParkerLab/danforth-2018.

### X-gal staining

X-gal experiments were conducted on *Gli1^tm2Alj^/J* (Jackson Laboratories, #008211) mice crossed to *Sd/+* mice. Embryos were dissected in PBS and fixed with Glutaraldehyde solution (1.25%: 1M EGTA, .2%: 1M MgCl_2_,2%: 25% Glutaraldehyde in PBS) and rinsed in wash buffer (.2%: 1MgCl_2_, .4%: 5% NP-40 in Sodium Phosphate buffer). Embryos were incubated in X-gal stain solution containing ferric and ferrocyanide ions for two hours. Embryos were then fixed with 4% PFA then rinsed and stored in PBS. (n ≥ 3 for all conditions) Images were taken with a Leica MZ10F dissecting microscope with a Fostec EKE ACE external light source.

### Whole mount in situ hybridization

Embryos were dissected in cold DEPC treated PBS, fixed in 4% paraformaldehyde (PFA) in DEPC treated PBS, washed in PBST, dehydrated through a graded methanol series (25%, 50%, 75%, 100%), and processed for in situ hybridization as described previously (81, 82). Embryos at each developmental stage for each probe were processed together, and images were captured under identical settings for comparison of staining intensity among WT, *Sd/+* and *Sd/Sd* genotypes, n ≥ 3 embryos of each genotype were studied with all probe conditions.

*Ptf1a* probes were generated from IMAGE clone: 8861527, MGC:170132, GenBank BC138507.1) in PCR4-TOPO. Sense probe was generated using digoxygenin labeling kit(Roche), T3 polymerase and Not I digested plasmid. Antisense probe was generated using T7 polymerase and Spe I digested plasmid. For the Gm13344 probe, a 511 bp fragment (Chr2: 19,351,134-19, 351,644) of Gm13344 lncRNA was amplified using primers GGGTGTATCACCCAGCAATC (Forward) and AAGAGGAGGAACCCAGGTGT (Reverse) from BAC DNA RP24-347M17, and cloned into PGEMT-easy vector. The antisense digoxygenin-AP probe was generated using SP6 polymerase from the Nco I digested PGEMT clone while the sense probe using T7 polymerase from Nde I digested clone. Probes for *Shh* (83–85), *Noto* (60), *Kirrel2/Neph3* (86), *Nkx 2-9* (40), *Foxa1, Foxa2* (87, 88), and *T* (89), were previously described.

### Immunofluorescence

Embryos were dissected into fresh, ice cold 4% PFA (Fluka: 76240) in phosphate buffered saline (PBS) for 30 minutes then cryoprotected overnight in solution of 10% sucrose and 2mM MgCl_2_ and cut immediately rostral to the forelimb and hindlimb buds. The tissues were embedded in OCT and stored at −80°. Frozen sections were taken through the transverse plane and stored at −80°. Tissue sections were blocked with PBS with 3%BSA, 1% Goat Serum, 0.1% Triton X-100, brought to a final pH of 7.4 for 1 hour at room temperature in a humidified container. Blocking solution was replaced with primary antibody diluted in blocking buffer. Primary antibodies were purchased from the Developmental Studies Hydronima Bank. SHH (5E1, AB_2188307), FOXA2 (4C7, AB_2278498), NKX2.2 (74.5A5, AB_2314952), NKX6.1(F55A10, AB_532378), PAX6 (PAX6, AB_528427). Tissue sections from n ≥ 3 embryos per genotype were incubated with each primary antibody overnight at 4° at the following concentrations. 1:20 for SHH, FOXA2, NKX2.2, NKX6.1, and 1:2500 for PAX6. Sections were stained with DAPI (Kirkegaard & Perry Laboratories: 71-03-00), incubated for one hour with Alexa fluor 488 goat anti-mouse secondary antibody (A11001), then coverslipped with aqueous mounting medium (Thermo Scientific: Immu-Mount: 990402) Fluorescent imaging was conducted on a Leitz DMRB microscope, with a Leica EL6000 fluorescent lamp.

### Caspase 3 Immunofluorescence

Embryos were processed for Paraffin embedding with a Tissue Tek VIP (Miles Scientific) and embedded in paraffin blocks. Blocks were sectioned with a Spencer Microtome (American Optical) at a thickness of 7μm and stored at room temperature. Paraffin was removed with Xylene followed by EtOH and PBS for rehydration. Antigen retrieval was achieved by citrate boiling and slides were then blocked with suppressor solution (5% goat serum, 3%BSA, 0.5% Tween 20) for 20 minutes. Sections (n > 3 per genotype) were incubated with Caspase 3 primary antibody (Cleaved Caspase-3 anti-rabbit, D175: 9661S Cell Signaling Technology) overnight at 4° at a concentration of 1:400. Sections were stained with DAPI (Kirkegaard & Perry Laboratories: 71-03-00), incubated for one hour with Alexa fluor 488 goat anti-rabbit secondary antibody (A11008), rinsed with PBS, then coverslips were applied with aqueous mounting medium (Thermo Scientific: Immu-Mount: 990402) Fluorescent imaging was conducted on a Leitz DMRB microscope, with a Leica EL6000 fluorescent lamp.

## Acknowledgements

The authors thank Drs. Sally Camper and Miriam Meisler for critical review of this manuscript. We acknowledge Dr. Andreas Kispert for the *Kirrel2/Neph3* probe, Dr. Hans-Henning Arnold for the *Nkx2-9* probe, Dr. Andreas Schedl for the *Nephrin/NPHS1* probe, Dr. Klaus Kaestner for *Foxa1* and *Foxa2* probes, and Dr. Cheryle Seguin for the *Noto* probe. We thank the University of Michigan DNA Sequencing Core for assistance with sequencing.

## Supporting information captions

**S1 Fig:**
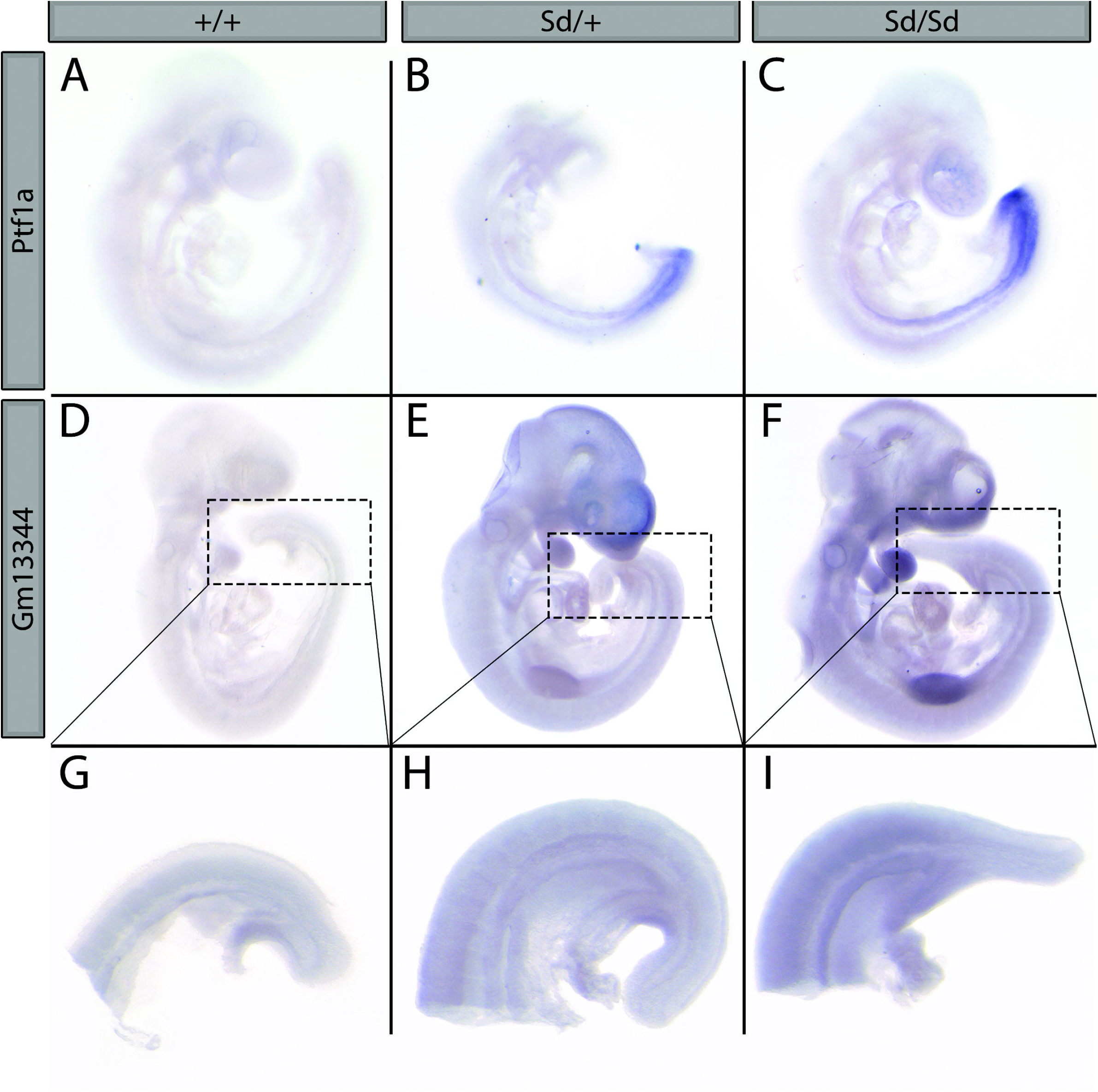
Whole mount In-Situ hybridization shows ectopic *Ptf1a* expression. (A-C) Whole mount in-situ hybridization with *Ptf1a* antisense probe in E9.5 WT, *Sd/+,* and *Sd/Sd* embryos. (D-I) Whole mount in situ hybridization with *Gm13344* antisense probe in E9.5 embryos and tailbuds.

S1 Table: **WT mouse tailbud transcriptome**. Expression values (FPKM) of genes in WT mouse tailbuds.

S2 Table: **Differential gene expression analysis results**. Results of the differential gene expression analysis using DESeq2. Positive fold change indicates higher expression in *Sd/Sd* mice.

S3 Table: **Somite counts for ATAC-seq and RNA-seq samples**. Each line in the table represents a single tailbud. If two rows share the same value in the ‘library’ column, then the two tailbuds were pooled as input for a single library. Because the number of somites was used as a covariate in some analyses, when two tailbuds were pooled the number of somites for the library was set to the average of the number of somites for the two individual embryos.

S4 Table: **Publicly available H3K4me1 ChIP-seq data analyzed**

## References

1 Pauli, R.M. (1994) Lower mesodermal defects: a common cause of fetal and early neonatal death. Am J Med Genet, 50, 154–172.

2 Dunn, L.C., Gluecksohn-Schoenheimer, S. and Bryson, V. (1940) A new mutation in the mouse affecting spinal column and urogenital system. Journal of Heredity, 31, 343–348.

3 Grüneberg, H. (1953) Genetioal studies on the skeleton of the mouse. Journal of Genetics, 51, 317–326.

4 Gluecksohn-Schoenheimer, S. (1945) The Embryonic Development of Mutants of the Sd-Strain in Mice. Genetics, 30, 29–38.

5 Vlangos, C.N., Siuniak, A.N., Robinson, D., Chinnaiyan, A.M., Lyons, R.H., Jr., Cavalcoli, J.D. and Keegan, C.E. (2013) Next-generation sequencing identifies the Danforth’s short tail mouse mutation as a retrotransposon insertion affecting Ptf1a expression. PLoS Genet, 9, e1003205.

6 Lugani, F., Arora, R., Papeta, N., Patel, A., Zheng, Z., Sterken, R., Singer, R.A., Caridi, G., Mendelsohn, C., Sussel, L. et al. (2013) A retrotransposon insertion in the 5ʹ regulatory domain of Ptf1a results in ectopic gene expression and multiple congenital defects in Danforth’s short tail mouse. PLoS Genet, 9, e1003206.

7 Semba, K., Araki, K., Matsumoto, K., Suda, H., Ando, T., Sei, A., Mizuta, H., Takagi, K., Nakahara, M., Muta, M. et al. (2013) Ectopic expression of Ptf1a induces spinal defects, urogenital defects, and anorectal malformations in Danforth’s short tail mice. PLoS Genet, 9, e1003204.

8 Krapp, A., Knofler, M., Frutiger, S., Hughes, G.J., Hagenbuchle, O. and Wellauer, P.K. (1996) The p48 DNA-binding subunit of transcription factor PTF1 is a new exocrine pancreas-specific basic helix-loop-helix protein. Embo j, 15, 4317–4329.

9 Willet, S.G., Hale, M.A., Grapin-Botton, A., Magnuson, M.A., MacDonald, R.J. and Wright, C.V. (2014) Dominant and context-specific control of endodermal organ allocation by Ptf1a. Development, 141, 4385–4394.

10 Obata, J., Yano, M., Mimura, H., Goto, T., Nakayama, R., Mibu, Y., Oka, C. and Kawaichi, M. (2001) p48 subunit of mouse PTF1 binds to RBP-Jkappa/CBF-1, the intracellular mediator of Notch signalling, and is expressed in the neural tube of early stage embryos. Genes Cells, 6, 345–360.

11 Glasgow, S.M., Henke, R.M., Macdonald, R.J., Wright, C.V. and Johnson, J.E. (2005) Ptf1a determines GABAergic over glutamatergic neuronal cell fate in the spinal cord dorsal horn. Development, 132, 5461–5469.

12 Sellick, G.S., Barker, K.T., Stolte-Dijkstra, I., Fleischmann, C., Coleman, R.J., Garrett, C., Gloyn, A.L., Edghill, E.L., Hattersley, A.T., Wellauer, P.K. et al. (2004) Mutations in PTF1A cause pancreatic and cerebellar agenesis. Nat Genet, 36, 1301–1305.

13 Krapp, A., Knofler, M., Ledermann, B., Burki, K., Berney, C., Zoerkler, N., Hagenbuchle, O. and Wellauer, P.K. (1998) The bHLH protein PTF1-p48 is essential for the formation of the exocrine and the correct spatial organization of the endocrine pancreas. Genes Dev, 12, 3752–3763.

14 Cockell, M., Stevenson, B.J., Strubin, M., Hagenbuchle, O. and Wellauer, P.K. (1989) Identification of a cell-specific DNA-binding activity that interacts with a transcriptional activator of genes expressed in the acinar pancreas. Mol Cell Biol, 9, 2464–2476.

15 Kawaguchi, Y., Cooper, B., Gannon, M., Ray, M., MacDonald, R.J. and Wright, C.V. (2002) The role of the transcriptional regulator Ptf1a in converting intestinal to pancreatic progenitors. Nat Genet, 32, 128–134.

16 Beres, T.M., Masui, T., Swift, G.H., Shi, L., Henke, R.M. and MacDonald, R.J. (2006) PTF1 is an organ-specific and Notch-independent basic helix-loop-helix complex containing the mammalian Suppressor of Hairless (RBP-J) or its paralogue, RBP-L. Mol Cell Biol, 26, 117–130.

17 Masui, T., Swift, G.H., Hale, M.A., Meredith, D.M., Johnson, J.E. and Macdonald, R.J. (2008) Transcriptional autoregulation controls pancreatic Ptf1a expression during development and adulthood. Mol Cell Biol, 28, 5458–5468.

18 Meredith, D.M., Masui, T., Swift, G.H., MacDonald, R.J. and Johnson, J.E. (2009) Multiple transcriptional mechanisms control Ptf1a levels during neural development including autoregulation by the PTF1-J complex. J Neurosci, 29, 11139–11148.

19 Mona, B., Avila, J.M., Meredith, D.M., Kollipara, R.K. and Johnson, J.E. (2016) Regulating the dorsal neural tube expression of Ptf1a through a distal 3ʹ enhancer. Dev Biol, 418, 216–225.

20 Buenrostro, J.D., Giresi, P.G., Zaba, L.C., Chang, H.Y. and Greenleaf, W.J. (2013) Transposition of native chromatin for fast and sensitive epigenomic profiling of open chromatin, DNA-binding proteins and nucleosome position. Nat Methods, 10, 1213–1218.

21 Scott, L.J., Erdos, M.R., Huyghe, J.R., Welch, R.P., Beck, A.T., Wolford, B.N., Chines, P.S., Didion, J.P., Narisu, N., Stringham, H.M. et al. (2016) The genetic regulatory signature of type 2 diabetes in human skeletal muscle. Nature communications, 7, 11764.

22 Varshney, A., Scott, L.J., Welch, R.P., Erdos, M.R., Chines, P.S., Narisu, N., Albanus, R.D., Orchard, P., Wolford, B.N., Kursawe, R. et al. (2017) Genetic regulatory signatures underlying islet gene expression and type 2 diabetes. Proc Natl Acad Sci U S A, 114, 2301–2306.

23 Lamparter, D., Marbach, D., Rueedi, R., Bergmann, S. and Kutalik, Z. (2017) Genome-Wide Association between Transcription Factor Expression and Chromatin Accessibility Reveals Regulators of Chromatin Accessibility. PLoS computational biology, 13, e1005311.

24 Cirillo, L.A., Lin, F.R., Cuesta, I., Friedman, D., Jarnik, M. and Zaret, K.S. (2002) Opening of compacted chromatin by early developmental transcription factors HNF3 (FoxA) and GATA-4. Mol Cell, 9, 279–289.

25 Chen, R. and Gifford, D.K. (2017) Differential chromatin profiles partially determine transcription factor binding. PloS one, 12, e0179411.

26 Sherwood, R.I., Hashimoto, T., O’Donnell, C.W., Lewis, S., Barkal, A.A., van Hoff, J.P., Karun, V., Jaakkola, T. and Gifford, D.K. (2014) Discovery of directional and nondirectional pioneer transcription factors by modeling DNase profile magnitude and shape. Nature biotechnology, 32, 171–178.

27 Love, M.I., Huber, W. and Anders, S. (2014) Moderated estimation of fold change and dispersion for RNA-seq data with DESeq2. Genome Biol, 15, 550.

28 Washietl, S., Kellis, M. and Garber, M. (2014) Evolutionary dynamics and tissue specificity of human long noncoding RNAs in six mammals. Genome Research, 24, 616–628.

29 Paralkar, V.R., Taborda, C.C., Huang, P., Yao, Y., Kossenkov, A.V., Prasad, R., Luan, J., Davies, J.O., Hughes, J.R., Hardison, R.C. et al. (2016) Unlinking an lncRNA from Its Associated cis Element. Mol Cell, 62, 104–110.

30 Weedon, M.N., Cebola, I., Patch, A.M., Flanagan, S.E., De Franco, E., Caswell, R., Rodriguez-Segui, S.A., Shaw-Smith, C., Cho, C.H., Allen, H.L. et al. (2014) Recessive mutations in a distal PTF1A enhancer cause isolated pancreatic agenesis. Nat Genet, 46, 61–64.

31 Heintzman, N.D., Stuart, R.K., Hon, G., Fu, Y., Ching, C.W., Hawkins, R.D., Barrera, L.O., Van Calcar, S., Qu, C., Ching, K.A. et al. (2007) Distinct and predictive chromatin signatures of transcriptional promoters and enhancers in the human genome. Nat Genet, 39, 311–318.

32 Kundaje, A., Meuleman, W., Ernst, J., Bilenky, M., Yen, A., Heravi-Moussavi, A., Kheradpour, P., Zhang, Z., Wang, J., Ziller, M.J. et al. (2015) Integrative analysis of 111 reference human epigenomes. Nature, 518, 317–330.

33 Thompson, N., Gesina, E., Scheinert, P., Bucher, P. and Grapin-Botton, A. (2012) RNA profiling and chromatin immunoprecipitation-sequencing reveal that PTF1a stabilizes pancreas progenitor identity via the control of MNX1/HLXB9 and a network of other transcription factors. Mol Cell Biol, 32, 1189–1199.

34 Meredith, D.M., Borromeo, M.D., Deering, T.G., Casey, B.H., Savage, T.K., Mayer, P.R., Hoang, C., Tung, K.C., Kumar, M., Shen, C. et al. (2013) Program specificity for Ptf1a in pancreas versus neural tube development correlates with distinct collaborating cofactors and chromatin accessibility. Mol Cell Biol, 33, 3166–3179.

35 O’Donnell, M., Hong, C.S., Huang, X., Delnicki, R.J. and Saint-Jeannet, J.P. (2006) Functional analysis of Sox8 during neural crest development in Xenopus. Development, 133, 3817–3826.

36 Abdelkhalek, H.B., Beckers, A., Schuster-Gossler, K., Pavlova, M.N., Burkhardt, H., Lickert, H., Rossant, J., Reinhardt, R., Schalkwyk, L.C., Muller, I. et al. (2004) The mouse homeobox gene Not is required for caudal notochord development and affected by the truncate mutation. Genes Dev, 18, 1725–1736.

37 Di Gregorio, A., Harland, R.M., Levine, M. and Casey, E.S. (2002) Tail morphogenesis in the ascidian, Ciona intestinalis, requires cooperation between notochord and muscle. Dev Biol, 244, 385–395.

38 Schulte-Merker, S., van Eeden, F.J., Halpern, M.E., Kimmel, C.B. and Nusslein-Volhard, C. (1994) no tail (ntl) is the zebrafish homologue of the mouse T (Brachyury) gene. Development, 120, 1009–1015.

39 Wilson, V., Manson, L., Skarnes, W.C. and Beddington, R.S. (1995) The T gene is necessary for normal mesodermal morphogenetic cell movements during gastrulation. Development, 121, 877–886.

40 Pabst, O., Herbrand, H. and Arnold, H.H. (1998) Nkx2-9 is a novel homeobox transcription factor which demarcates ventral domains in the developing mouse CNS. Mech Dev, 73, 85–93.

41 Pabst, O., Herbrand, H., Takuma, N. and Arnold, H.H. (2000) NKX2 gene expression in neuroectoderm but not in mesendodermally derived structures depends on sonic hedgehog in mouse embryos. Development genes and evolution, 210, 47–50.

42 Weinstein, D.C., Ruiz i Altaba, A., Chen, W.S., Hoodless, P., Prezioso, V.R., Jessell, T.M. and Darnell, J.E., Jr. (1994) The winged-helix transcription factor HNF-3 beta is required for notochord development in the mouse embryo. Cell, 78, 575–588.

43 Epstein, D.J., McMahon, A.P. and Joyner, A.L. (1999) Regionalization of Sonic hedgehog transcription along the anteroposterior axis of the mouse central nervous system is regulated by Hnf3-dependent and -independent mechanisms. Development, 126, 281–292.

44 Jeong, Y. and Epstein, D.J. (2003) Distinct regulators of Shh transcription in the floor plate and notochord indicate separate origins for these tissues in the mouse node. Development, 130, 3891–3902.

45 Litingtung, Y. and Chiang, C. (2000) Specification of ventral neuron types is mediated by an antagonistic interaction between Shh and Gli3. Nature neuroscience, 3, 979–985.

46 Ribes, V., Balaskas, N., Sasai, N., Cruz, C., Dessaud, E., Cayuso, J., Tozer, S., Yang, L.L., Novitch, B., Marti, E. et al. (2010) Distinct Sonic Hedgehog signaling dynamics specify floor plate and ventral neuronal progenitors in the vertebrate neural tube. Genes Dev, 24, 1186–1200.

47 Briscoe, J., Sussel, L., Serup, P., Hartigan-O’Connor, D., Jessell, T.M., Rubenstein, J.L. and Ericson, J. (1999) Homeobox gene Nkx2.2 and specification of neuronal identity by graded Sonic hedgehog signalling. Nature, 398, 622–627.

48 Noguchi, K.K., Cabrera, O.H., Swiney, B.S., Salinas-Contreras, P., Smith, J.K. and Farber, N.B. (2015) Hedgehog regulates cerebellar progenitor cell and medulloblastoma apoptosis. Neurobiology of disease, 83, 35–43.

49 Sharma, N., Nanta, R., Sharma, J., Gunewardena, S., Singh, K.P., Shankar, S. and Srivastava, R.K. (2015) PI3K/AKT/mTOR and sonic hedgehog pathways cooperate together to inhibit human pancreatic cancer stem cell characteristics and tumor growth. Oncotarget, 6, 32039–32060.

50 Wang, X., Wei, S., Zhao, Y., Shi, C., Liu, P., Zhang, C., Lei, Y., Zhang, B., Bai, B., Huang, Y. et al. (2017) Anti-proliferation of breast cancer cells with itraconazole: Hedgehog pathway inhibition induces apoptosis and autophagic cell death. Cancer Lett, 385, 128–136.

51 Zhao, W., Pan, X., Li, T., Zhang, C. and Shi, N. (2016) Lycium barbarum Polysaccharides Protect against Trimethyltin Chloride-Induced Apoptosis via Sonic Hedgehog and PI3K/Akt Signaling Pathways in Mouse Neuro-2a Cells. Oxidative medicine and cellular longevity, 2016, 9826726.

52 Chen, K.Y., Chiu, C.H. and Wang, L.C. (2017) Anti-apoptotic effects of Sonic hedgehog signalling through oxidative stress reduction in astrocytes co-cultured with excretory-secretory products of larval Angiostrongylus cantonensis. Scientific reports, 7, 41574.

53 Latos, P.A., Pauler, F.M., Koerner, M.V., Senergin, H.B., Hudson, Q.J., Stocsits, R.R., Allhoff, W., Stricker, S.H., Klement, R.M., Warczok, K.E. et al. (2012) Airn transcriptional overlap, but not its lncRNA products, induces imprinted Igf2r silencing. Science, 338, 1469–1472.

54 Ristola, M. and Lehtonen, S. (2014) Functions of the podocyte proteins nephrin and Neph3 and the transcriptional regulation of their genes. Clinical science (London, England : 1979), 126, 315–328.

55 Nishida, K., Hoshino, M., Kawaguchi, Y. and Murakami, F. (2010) Ptf1a directly controls expression of immunoglobulin superfamily molecules Nephrin and Neph3 in the developing central nervous system. J Biol Chem, 285, 373–380.

56 Jessell, T.M. (2000) Neuronal specification in the spinal cord: inductive signals and transcriptional codes. Nat Rev Genet, 1, 20–29.

57 Herrmann, B.G., Labeit, S., Poustka, A., King, T.R. and Lehrach, H. (1990) Cloning of the T gene required in mesoderm formation in the mouse. Nature, 343, 617–622.

58 Pennimpede, T., Proske, J., Konig, A., Vidigal, J.A., Morkel, M., Bramsen, J.B., Herrmann, B.G. and Wittler, L. (2012) In vivo knockdown of Brachyury results in skeletal defects and urorectal malformations resembling caudal regression syndrome. Dev Biol, 372, 55–67.

59 Plouhinec, J.L., Granier, C., Le Mentec, C., Lawson, K.A., Saberan-Djoneidi, D., Aghion, J., Shi, D.L., Collignon, J. and Mazan, S. (2004) Identification of the mammalian Not gene via a phylogenomic approach. Gene expression patterns : GEP, 5, 11–22.

60 McCann, M.R., Tamplin, O.J., Rossant, J. and Seguin, C.A. (2012) Tracing notochord-derived cells using a Noto-cre mouse: implications for intervertebral disc development. Disease models & mechanisms, 5, 73–82.

61 Ang, S.L. and Rossant, J. (1994) HNF-3 beta is essential for node and notochord formation in mouse development. Cell, 78, 561–574.

62 Chiang C, L.Y., Lee E, Young KE, Corden JL, Westphal H, Beachy PA. (1996) Cyclopia and defective axial patterning in mice lacking Sonic hedgehog gene function. Nature, 383, 407–413.

63 Mo, R., Kim, J.H., Zhang, J., Chiang, C., Hui, C.C. and Kim, P.C. (2001) Anorectal malformations caused by defects in sonic hedgehog signaling. Am J Pathol, 159, 765–774.

64 Kimmel, S.G., Mo, R., Hui, C.C. and Kim, P.C. (2000) New mouse models of congenital anorectal malformations. J Pediatr Surg, 35, 227–230; discussion 230-221.

65 Kim, P.C.W., Mo, R. and Hui, C.-c. (2001) Murine models of VACTERL syndrome: Role of sonic hedgehog signaling pathway. Journal of Pediatric Surgery, 36, 381–384.

66 Runck, L.A., Method, A., Bischoff, A., Levitt, M., Pena, A., Collins, M.H., Gupta, A., Shanmukhappa, S., Wells, J.M. and Guasch, G. (2014) Defining the molecular pathologies in cloaca malformation: similarities between mouse and human. Disease models & mechanisms, 7, 483–493.

67 Choi, K.-S., Lee, C. and Harfe, B.D. (2012) Sonic hedgehog in the notochord is sufficient for patterning of the intervertebral discs. Mechanisms of development, 129, 255–262.

68 Maier, J.A., Lo, Y. and Harfe, B.D. (2013) Foxa1 and Foxa2 Are Required for Formation of the Intervertebral Discs. PloS one, 8, e55528.

69 Picelli, S., Bjorklund, A.K., Reinius, B., Sagasser, S., Winberg, G. and Sandberg, R. (2014) Tn5 transposase and tagmentation procedures for massively scaled sequencing projects. Genome Res, 24, 2033–2040.

70 Li, H. and Durbin, R. (2009) Fast and accurate short read alignment with Burrows-Wheeler transform. Bioinformatics, 25, 1754–1760.

71 Li, H., Handsaker, B., Wysoker, A., Fennell, T., Ruan, J., Homer, N., Marth, G., Abecasis, G., Durbin, R. and Genome Project Data Processing, S. (2009) The Sequence Alignment/Map format and SAMtools. Bioinformatics, 25, 2078–2079.

72 Zhang, Y., Liu, T., Meyer, C.A., Eeckhoute, J., Johnson, D.S., Bernstein, B.E., Nusbaum, C., Myers, R.M., Brown, M., Li, W. et al. (2008) Model-based analysis of ChIP-Seq (MACS). Genome Biol, 9, R137.

73 Quinlan, A.R. (2014) BEDTools: The Swiss-Army Tool for Genome Feature Analysis. Current protocols in bioinformatics, 47, 11.12.11–34.

74 Aken, B.L., Ayling, S., Barrell, D., Clarke, L., Curwen, V., Fairley, S., Fernandez Banet, J., Billis, K., García Girón, C., Hourlier, T. et al. (2016) The Ensembl gene annotation system. Database, 2016, baw093–baw093.

75 Djebali, S., Davis, C.A., Merkel, A., Dobin, A., Lassmann, T., Mortazavi, A., Tanzer, A., Lagarde, J., Lin, W., Schlesinger, F. et al. (2012) Landscape of transcription in human cells. Nature, 489, 101–108.

76 Hartley, S.W. and Mullikin, J.C. (2015) QoRTs: a comprehensive toolset for quality control and data processing of RNA-Seq experiments. BMC bioinformatics, 16, 224.

77 Lee, C., Patil, S. and Sartor, M.A. (2016) RNA-Enrich: a cut-off free functional enrichment testing method for RNA-seq with improved detection power. Bioinformatics, 32, 1100–1102.

78 Landt, S.G., Marinov, G.K., Kundaje, A., Kheradpour, P., Pauli, F., Batzoglou, S., Bernstein, B.E., Bickel, P., Brown, J.B., Cayting, P. et al. (2012) ChIP-seq guidelines and practices of the ENCODE and modENCODE consortia. Genome Res, 22, 1813–1831.

79 Kharchenko, P.V., Tolstorukov, M.Y. and Park, P.J. (2008) Design and analysis of ChIP-seq experiments for DNA-binding proteins. Nature biotechnology, 26, 1351–1359.

80 Denas, O., Sandstrom, R., Cheng, Y., Beal, K., Herrero, J., Hardison, R.C. and Taylor, J. (2015) Genome-wide comparative analysis reveals human-mouse regulatory landscape and evolution. BMC Genomics, 16, 87.

81 Keegan, C.E., Hutz, J.E., Else, T., Adamska, M., Shah, S.P., Kent, A.E., Howes, J.M., Beamer, W.G. and Hammer, G.D. (2005) Urogenital and caudal dysgenesis in adrenocortical dysplasia (acd) mice is caused by a splicing mutation in a novel telomeric regulator. Hum Mol Genet, 14, 113–123.

82 Vlangos, C.N., O’Connor, B.C., Morley, M.J., Krause, A.S., Osawa, G.A. and Keegan, C.E. (2009) Caudal regression in adrenocortical dysplasia (acd) mice is caused by telomere dysfunction with subsequent p53-dependent apoptosis. Dev Biol, 334, 418–428.

83 Harfe, B.D., Scherz, P.J., Nissim, S., Tian, H., McMahon, A.P. and Tabin, C.J. (2004) Evidence for an expansion-based temporal Shh gradient in specifying vertebrate digit identities. Cell, 118, 517–528.

84 Echelard, Y., Epstein, D.J., St-Jacques, B., Shen, L., Mohler, J., McMahon, J.A. and McMahon, A.P. (1993) Sonic hedgehog, a member of a family of putative signaling molecules, is implicated in the regulation of CNS polarity. Cell, 75, 1417–1430.

85 Everson, J.L., Fink, D.M., Yoon, J.W., Leslie, E.J., Kietzman, H.W., Ansen-Wilson, L.J., Chung, H.M., Walterhouse, D.O., Marazita, M.L. and Lipinski, R.J. (2017) Sonic hedgehog regulation of Foxf2 promotes cranial neural crest mesenchyme proliferation and is disrupted in cleft lip morphogenesis. Development, 144, 2082–2091.

86 Volker, L.A., Petry, M., Abdelsabour-Khalaf, M., Schweizer, H., Yusuf, F., Busch, T., Schermer, B., Benzing, T., Brand-Saberi, B., Kretz, O. et al. (2012) Comparative analysis of Neph gene expression in mouse and chicken development. Histochemistry and cell biology, 137, 355–366.

87 Monaghan, A.P., Kaestner, K.H., Grau, E. and Schutz, G. (1993) Postimplantation expression patterns indicate a role for the mouse forkhead/HNF-3 alpha, beta and gamma genes in determination of the definitive endoderm, chordamesoderm and neuroectoderm. Development, 119, 567–578.

88 Kaestner, K.H., Hiemisch, H., Luckow, B. and Schütz, G. (1994) The HNF-3 Gene Family of Transcription Factors in Mice: Gene Structure, cDNA Sequence, and mRNA Distribution. Genomics, 20, 377–385.

89 Wilson, V., Rashbass, P. and Beddington, R.S. (1993) Chimeric analysis of T (Brachyury) gene function. Development, 117, 1321–1331.

